# Accurate quantification of canine mitochondrial DNA copy number and its evaluation as a biomarker of brain injury

**DOI:** 10.64898/2026.02.16.706235

**Authors:** Eliane Caseiro Soares de Menezes, Karolina Sek, Abbe Crawford, Minjie Zhang, Anqi Shi, Nimrah Rauf, Claire Thornton, Afshan N Malik

## Abstract

Acute brain injury is challenging to manage in veterinary medicine, with limited validated means of prognostication currently available. The brain is rich in mitochondrial content and contains thousands of copies of mitochondrial DNA (mtDNA) per cell. We hypothesized that brain cell loss following acute brain injury may result in release of mtDNA into the systemic circulation. To investigate this, mtDNA-CN was measured in blood (n=4-6/group) and cerebral cortex (n=1/group) samples from dogs with and without brain injury using absolute quantification by real-time qPCR. By filtering out regions with homology to nuclear mitochondrial insertion sequences (NumtS) and repetitive regions, oligonucleotide primers were designed to the canine mitochondrial and nuclear genomes. In controls, blood mtDNA-CN ranged from 98 to 288 (mean 193±72), in cases of brain injury, there was a non-significant trend for higher mtDNA-CN, ranging from 163-453 (mean 244±106). As expected, cerebral cortex contained higher mtDNA-CN than the blood. In a single case with serial sampling, mtDNA-CN increased five days post-injury. We present for the first time an assay to accurately quantify mtDNA-CN in canine samples, with potential as a biomarker for acute brain injury in veterinary practice.

**Simple Summary:** This work describes a novel assay to accurately measure absolute levels of canine mitochondrial DNA copy number (mtDNA-CN) in biological samples. We describe the range of mtDNA-CN in canine blood and show pilot data suggesting that blood mtDNA-CN should be evaluated as a potential biomarker of acute brain injury.

## 1. Introduction

Acute brain injury can cause severe neurological deficits and is associated with high morbidity and mortality [1]. In domesticated canines, common causes include road traffic accidents, other traumatic injuries, infections, immune mediated inflammation, toxicity, and seizures [2]. Primary injury to the central nervous system (CNS) can result in alteration of cellular homeostasis, neuroinflammation and neuronal death [3]. In humans, advanced imaging tools, such as computed tomography (CT), or magnetic resonance imaging (MRI) are used for the detection of and prognostication in CNS injury [4]. However, the use of neuroimaging tools in veterinary practice is limited due to high costs and the need for sedation or general anaesthesia in potentially unstable animals. Medical and surgical therapeutic options are available for dogs with brain injury, but there is a need for validated biomarkers of brain injury which could assist in rapid diagnosis and accurate prognostication.

Located in the cytosol of eukaryotic cells, and generating ATP to support cellular energy requirements, mitochondria contain a circular DNA genome (mtDNA), present in hundreds to thousands of copies per cell, depending on the cells’ bioenergetic requirements [5]. Believed to have been acquired by a host cell engulfing a prokaryotic cell via symbiosis during the evolution of the eukaryotic cell, mtDNA resembles bacterial genomes in several ways: it is a highly conserved circular double stranded DNA molecule of 16.6 kb in humans, and ∼16.7 kb in canines, which has a similar methylation pattern, replication and transcription processes to prokaryotes [6-8]. MtDNA is located embedded in the inner mitochondrial membrane complexed with the protein TFAM, and being close to the electron transport chain, it is more prone to damage than nuclear DNA, which is well protected through being packaged within histone proteins and located in the nucleus, a separate compartment of the cell [9]. mtDNA is often present in multiple copies per mitochondrion, while the number of mitochondria per cell is also highly variable between different cell types and can change in response to physiological signals including acute injury [10]. In the normal cell, damaged mtDNA is enzymatically degraded during the normal mitochondrial life cycle. However, in conditions of disease or acute injury, mtDNA can leak into the periphery where, because of its resemblance to bacterial DNA, it can activate the TLR-9 pathway, leading to TNF-alpha synthesis and a downstream inflammatory response [11].

We proposed the use of mtDNA-CN as a biomarker of mitochondrial dysfunction originally in 2011[10, 12]. Since then, changes in human mtDNA-CN in body fluids have been widely reported [13] including studies showing that circulating mtDNA-CN correlates with acute injury and is a predictor of mortality in patients in ICU [14, 15]. Injury in tissues rich in mitochondria, such as the brain, heart or kidneys, can result in release of mitochondria into the periphery and there is increasing evidence that this can result in subsequent inflammation [16, 17]. The brain has a very high mtDNA content, with regions of the human brain containing hundreds of thousands of copies of mtDNA per cell [18]. Therefore, we speculated that brain injury may result in the release of mtDNA into the periphery. Mitochondrial dysfunction has been shown to contribute to cell death following brain injury [19] and mtDNA levels were shown to change in a porcine model of traumatic brain injury [20]. However, to date, there has been no studies investigating the role of canine mtDNA-CN in brain injury.

A common method for measuring mtDNA content is to quantify a mitochondrial encoded gene relative to a nuclear encoded gene to determine the mitochondrial genome to nuclear genome ratio which we have termed Mt/N[12]. The exact copy numbers of mtDNA and nuclear DNA present in the sample can be measured using either absolute quantification or digital PCR [21, 22]. Both require small amounts of sample, are highly sensitive methods and can be easily adapted for high throughput clinical use [23]. When designing primers to use for Mt/N determination, it is important to identify and avoid nuclear mitochondrial insertion sequences (NumtS), regions of mtDNA which show high similarity to the nuclear genome, in order to avoid their co-amplification [12, 24]. As NumtS have been previously described in the canine genome [25], any assay for measuring canine mtDNA needs to avoid measuring these regions.

In the current study, our aim was to design an assay for measuring canine mtDNA-CN from biological samples. To do this we set up a real time qPCR assay by designing primers for the mitochondrial genome which do not co-amplify NumtS, and primers for the nuclear genome which are in a stable non-duplicated region. Although mtDNA-CN has recently been reported using relative qPCR approaches in the peripheral blood of dogs with heart failure [26] and in ageing castrated dogs [27], here we report absolute quantification, enabling direct comparisons between tissues. We measured mtDNA-CN in blood and cerebral cortex samples from dogs with and without brain injury to describe tissue-specific levels and to explore its potential as a biomarker of brain injury in veterinary medicine.

## 2. Materials and Methods

### 2.1. Animals

Residual blood samples were collected at the Royal Veterinary College’s (RVC) Queen Mother Hospital for Animals. Brain tissue was collected at post-mortem examination performed at the RVC’s Anatomical Pathology Department. Approval for this work was obtained from the local Ethical Review Board. Baseline characteristics and reason for euthanasia can be found in Table 1.

**Table 1.**
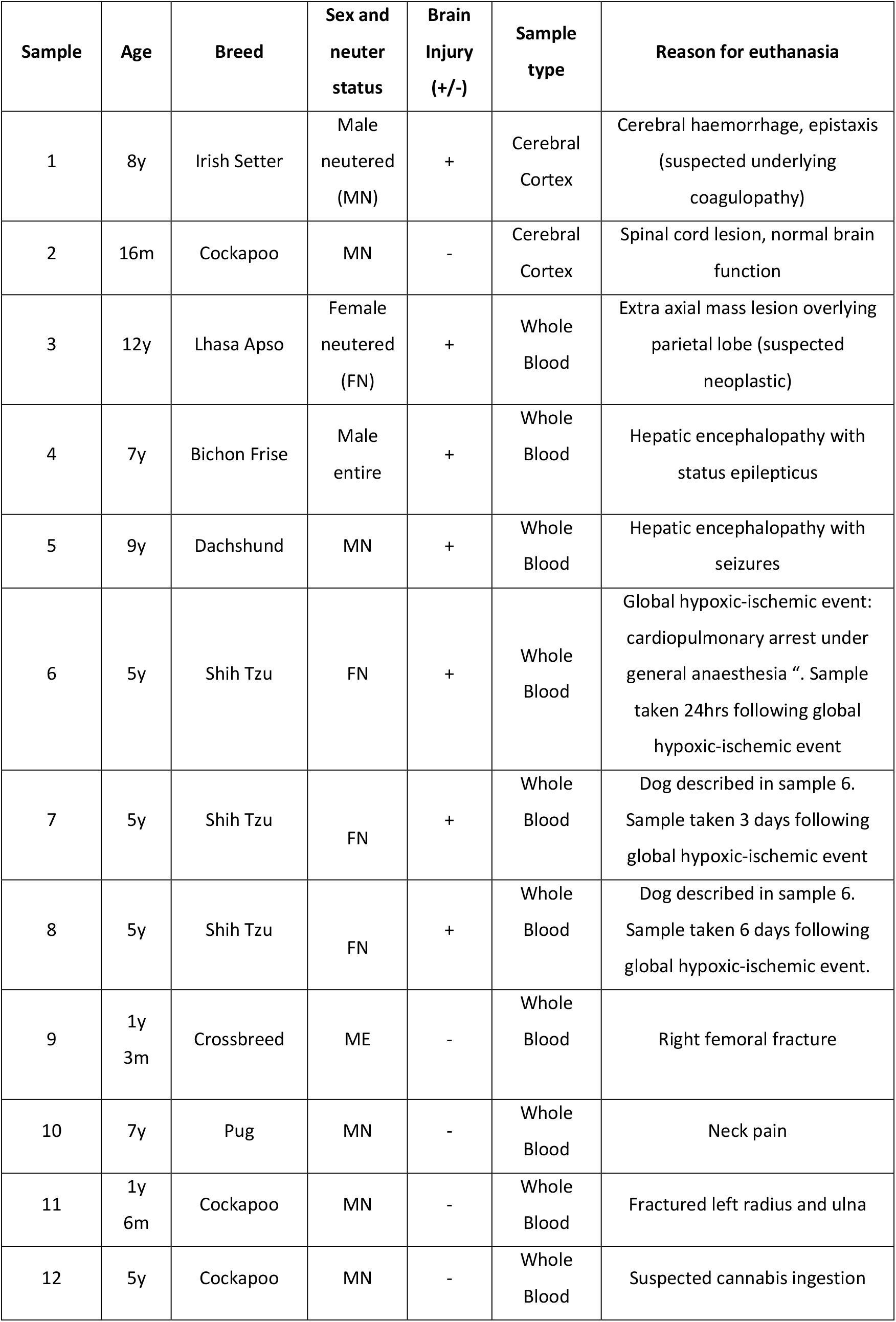
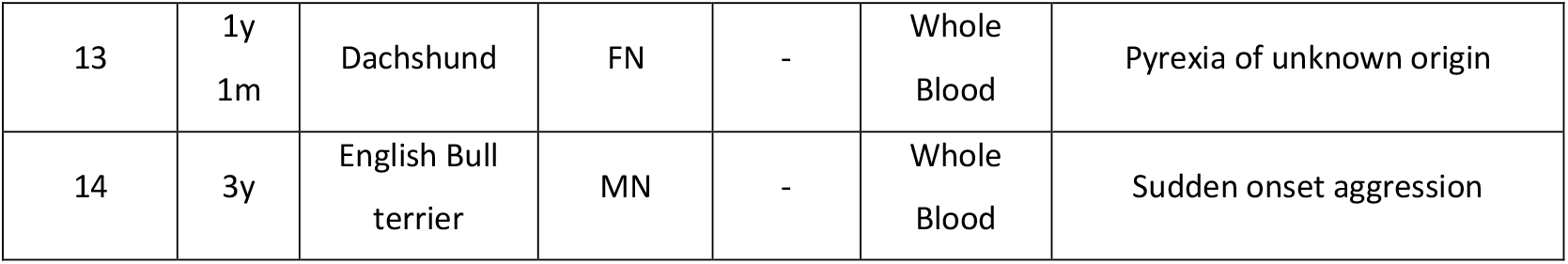
Information of canine biological samples used in this study. Table containing baseline characteristics of canine samples used in this study including age (y=years, m= months), brain injury (+= yes, −=no), sample type and reason for euthanasia.

### 2.2. Genomic DNA preparation

DNA isolation was performed using DNeasy Blood and Tissue Kit (Qiagen, UK) following the manufacturer’s instructions. DNA yield and quality were assessed using NanoDrop. The concentration of the DNA was adjusted to 10 ng/µl using nuclease-free water. To minimize the dilution bias effect and to improve the accuracy of qPCR analysis, genomic DNA was subjected to a 10-minute sonification using a bath sonication with a frequency of 38 kHz (Kerry, Pulsa-Tron 55) as previously described [12]. To avoid errors arising from repeated freeze thaw cycles, DNA samples were stored at 4°C for the duration of the study.

### 2.3. Identification of nuclear mitochondrial insertion sequences in the canine nuclear genome and identification of unique regions

The presence and location of NumtS within the canine nuclear genome was identified using BLAST (https://www.ncbi.nlm.nih.gov). Unique mtDNA regions with no homology to NumtS were identified by comparison of the mitochondrial genome of two different breeds of domestic dogs (Accession numbers: NC_002008.4 and CM025140.1) against the representative canine nuclear genome (taxid: 9615). The location of NumtS within the canine nuclear genome was examined using the BLAST alignment inspector tool as part of the NCBI genome data viewer. Regions in mtDNA with no or low similarity scores to the NumtS within chromosomal DNA of the given species were selected as suitable for primer design.

### 2.4. Development of accurate assays for absolute quantification of canine mtDNA-CN

Primers were designed to the unique mitochondrial region (forward, cMitoF1; reverse, cMitoR1) and single copy nuclear gene, beta-2 microglobulin (B2M) (forward, cB2MF1; reverse, cB2MR1) using BLAST (https://www.ncbi.nlm.nih.gov/tools/primer-blast) software (Figure 1). Primers were synthesized at Integrated DNA Technologies, UK.

**Figure 1.**
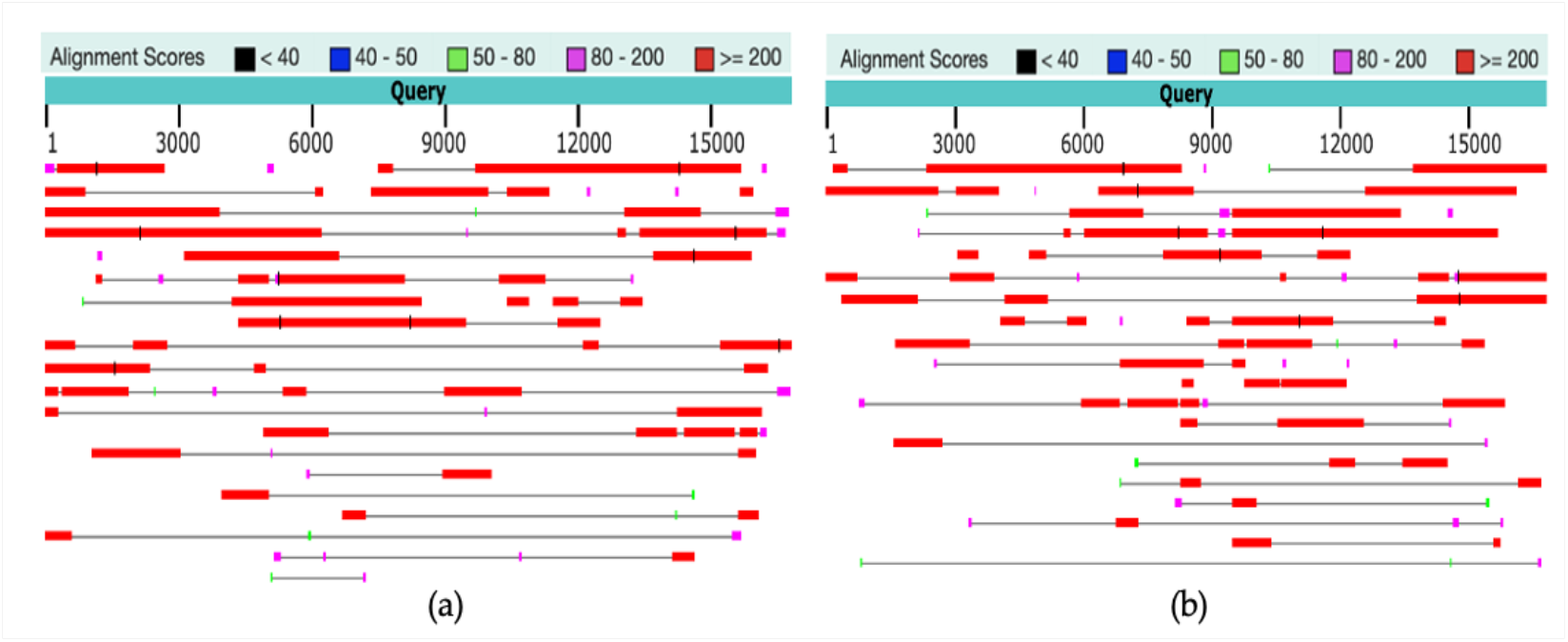
High sequence identity (<95%) is present in canine mitochondrial to the nuclear genome. (a) Analysis was performed by blasting canine reference genome against mitochondrial genomic sequences of two canine breeds; (a) Boxer (accession no. NC_002008.4); and (b) Golden Retriever (accession no. CM025140.1). Top hits are represented as coloured bar scattered at genomic locations; the colour key foralignment score is provided at the top of the figure; with red being the highest and black being the lowest score.

PCR amplicons from canine mtDNA (cMitoF1-R1) and canine B2M (cB2MF1-R1) primers were purified and used to prepare dilution standards for qPCR to allow for absolute qPCR quantification as follows: DNA products were excised from 1% agarose gel electrophoresis and purified using QIAquick gel extraction kit following the manufacturer’s instructions (Qiagen, UK). Concentration was measured using NanoDrop and DNA copy number in the stock solution was calculated as previously described [20]. The dilution range of 10^2^-10^8^ copies per 2 µL was prepared by diluting stock solution (10 µg/ml) with nuclease-free water (Sigma, UK). Purity of standards was evaluated by the presence of a single band when electrophoresed on a 1% agarose gel. All samples were stored at 4°C for the duration of the study.

MtDNA-CN from cases and controls were determined from template DNA by carrying out qPCR in a total volume of 10 µL containing 5 µL of QuantiFast SYBR Master Mix (Qiagen, 204054), 2 µL of nuclease-free water, 0.5 µL of forward and reverse primers (400nM final concentration each) for MtDNA (cMitoF1-R1) and B2M (cB2MF1-R1), and 2 µL of pre-treated genomic DNA. The reactions were performed in a LightCycler 480 (LC, Roche) using the following protocol: pre-incubation at 5 min at 9°C (1 cycle); denaturation for 10s at 95°C and annealing for 30s at 60°C (40 cycles); melting for 5s at 96°C and 60s at 65°C (1 cycle; for melting curve analysis); cooling for 30s at 4°C. All samples were run in triplicate. The standard curve was generated, and primer specificity was confirmed by a single melt peak at 82°C for cMito and 82°C for cB2M using LightCycler 96 software. qPCR efficiency was calculated from the slope between 95% and 105% with co-efficiency of reaction R^2^=0.98-0.99. To calculate mtDNA-CN, the mean value of the mitochondrial gene (Mito) was divided by the mean value of a nuclear gene (B2M) and then doubled to account for diploid nucleus. MtDNA-CN was expressed as Mt/N ratio.

## 3. Results

### 3.1. Identification of nuclear mitochondrial insertion sequences (NumtS) in the canine nuclear genome and identification of unique regions in the mitochondrial genome

The canine mitochondrial genomic sequence (accession numbers; NC_002008.4 and CM025140.1) was derived from two different breeds belonging to Canis lupus familiars and was searched against the canine reference genome (RefSeq, taxid:9615) using BLAST (https://www.ncbi.nlm.nih.gov/tools/primer-blast). The resulting data showed high sequence identity (<95%) of canine mitochondrial to nuclear genome suggesting the presence of NumtS. These were scattered at different positions within the nuclear genome and were present in 30 out of 39 chromosomes accounting for ∼1.2 % of the genome. The NumtS were present at various lengths ranging from 34-3944 bp with the largest being identical to ∼36% of the entire mtDNA sequence (Figure 1).

The majority of NumtS (∼57%) were positioned within the intergenic regions of the genome, compared to around (∼43%) of NumtS located within genes. Further inspection of NUMT position within genes revealed only 2 NumtS were located within the exonic region of the gene; these were positioned in exon 13 of the AGFG2 gene on chromosome 6 and exon 4 of an uncharacterized locus occupying chromosome 7. The remaining NumtS located within genes were found within the intronic region of the genes with a total of 15 NumtS occupying uncharacterized loci within the canine genome.

Canine mitochondrial genome was analysed further with BLAST and Primer-BLAST (https://www.ncbi.nlm.nih.gov); mitochondrial sequences were split into 150bp fragments (with 50bp overlaps), and each was blasted against the reference genome to search for unique mitochondrial fragments. Mitochondrial genome sequences from two breeds with dissimilarities between their mitochondrial genomes were used to enhance the process of finding and designing primers that do not co-amplify NumtS. A unique sequence that matched both Boxer and Golden Retriever breeds was identified at the position 14,231-14,403 and 6,927-7,099 bp of their mitochondrial genomes respectively. This sequence was used to design mitochondrial primers (forward, cMitoF1; reverse, cMitoR1) whereas genomic primers (forward, cB2MF1; reverse, cB2MR1) were designed against the B2M gene of the nuclear genome (Table 2).

**Table 2.**
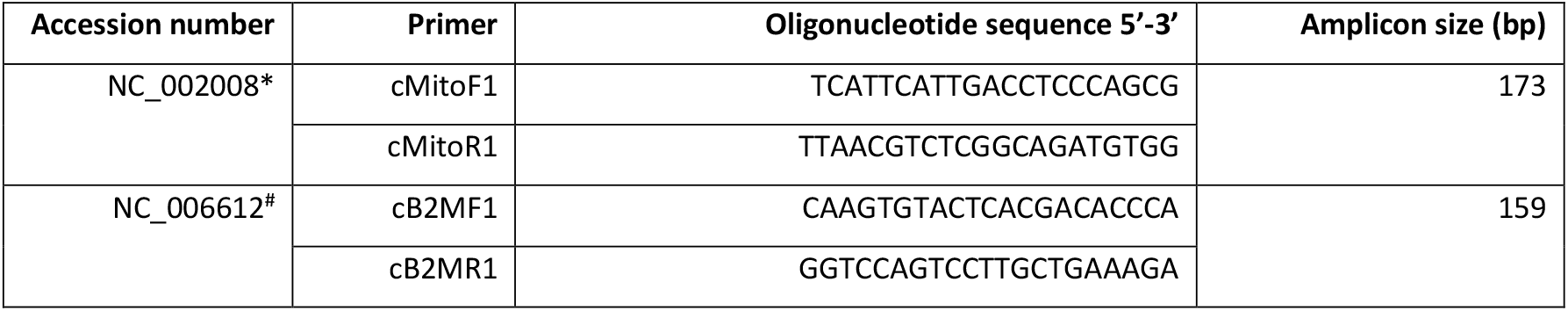
Canine primers sequences, accession number and amplicon size. ^*^The primer set cMitoF1-cMitoR1 can also work for Accession number CM025140. ^*#*^The primer set cB2MF1-cB2MR1 can also work for Accession number NC_051834.

### 3.2 Assay for absolute quantification of mtDNA-CN

To test the reproducibility and validity of the assay, primers were initially tested using standard PCR; the resulting products visualized with gel electrophoresis showed successful amplification of expected size products without visible co-amplification of other products. Absolute quantification of mtDNA-CN was performed using real-time qPCR; as previously described [28]. The target samples were run in triplicate using canine primers (cMitoF1-R1, cB2MF1-R1) and standards were run in quadruples. The standard curve obtained from 10-fold serial dilution samples was shown to be linear with a R2 value of 0.99 for both mitochondrial and nuclear DNA with an amplification efficiency of 97% and 95% respectively (Figure 2 and Figure 3). Both melting curves contained a single peak further confirming the specificity of the primer (Figure 2C and 3C).

**Figure 2.**
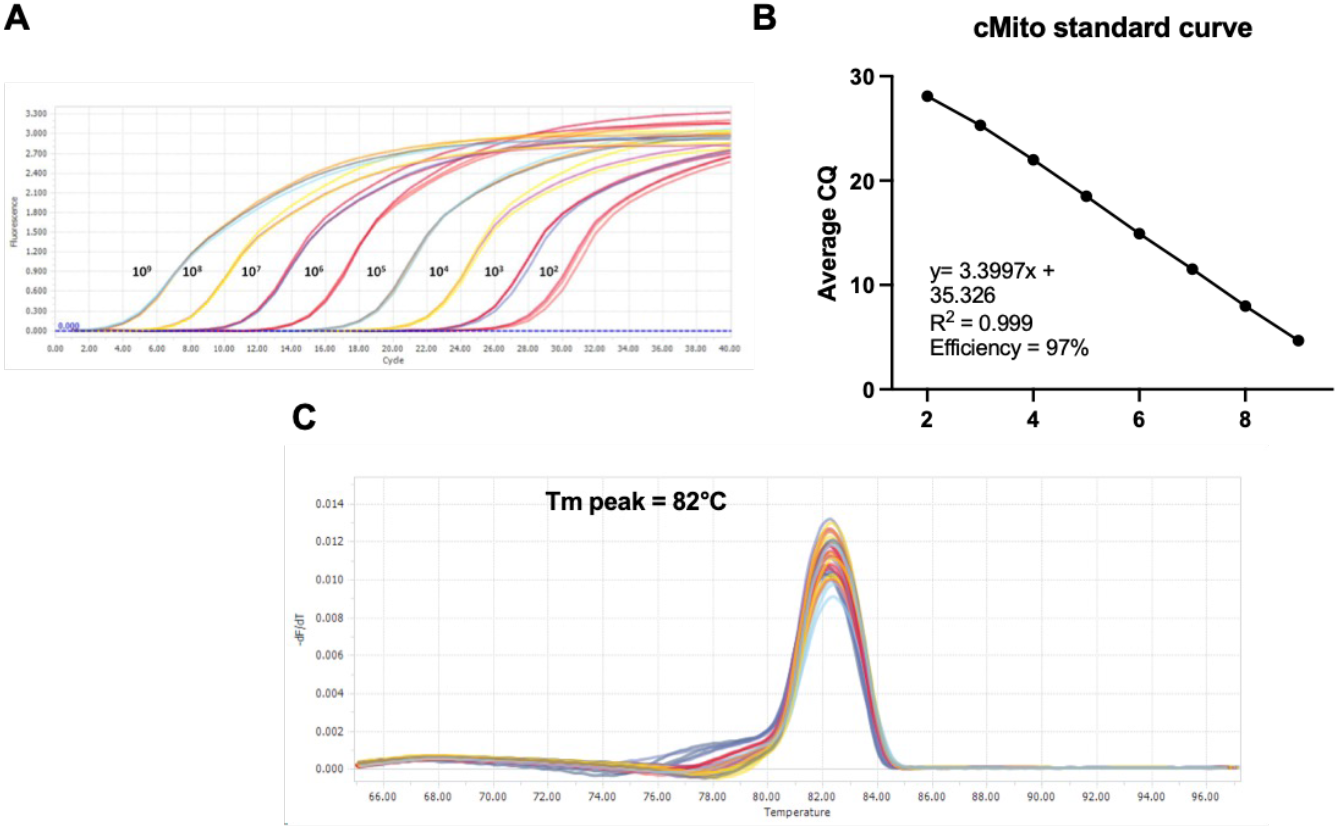
Amplification of cMT to generate a standard curve and Tm analysis. Dilution standards of cMito were prepared and 2µL of each standard was used in qPCR reaction and amplified. **(A)** Fluorescence data was acquired once per cycle and amplification curve was generated showing dilution series from 10^9^ to 10^2^ copies of cMito **(B)** CQ values for dilution standards were plotted against log copy number to generate standard curve. **(C)** Melting point analysis indicates specificity of amplified product as one single melt peak is visible at 82°C.

**Figure 3.**
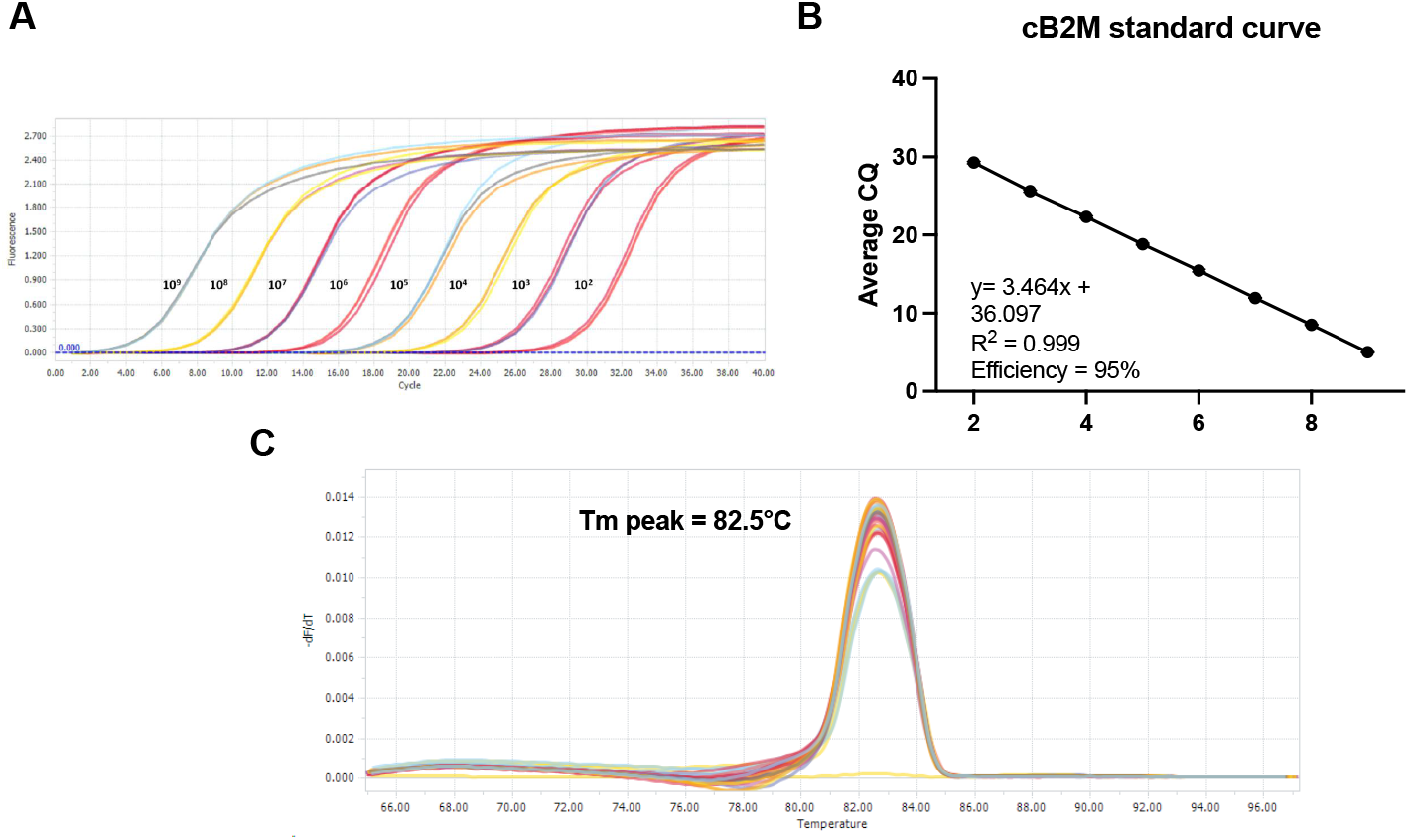
Amplification of cB2M to generate a standard curve and Tm analysis. Dilution standards of cB2M were prepared and 2µL of each standard was used in qPCR reaction and amplified. (**A**). Fluorescence data was acquired once per cycle and amplification curve was generated showing dilution series from 10^9^ to 10^2^ copies of cB2M **(B)** CQ values for dilution standards were plotted against log copy number to generate standard curve. **(C)** Melting point analysis indicates specificity of amplified product as one single melt peak is visible at 82.5°C.

### 3.4 Blood and cerebral cortex mtDNA-CN in dogs with and without suspected brain injury

Blood MtDNA-CN in dogs with no brain injury ranged from 98 to 288 copies/genome with a mean of 193 ±72. In dogs with brain injury, the blood mtDNA-CN spanned a larger range between 163 and 453 with a mean of 244 ±106 (Table 3).

**Table 3.** Absolute quantification of mitochondrial DNA and nuclear DNA: comparison of data from brain injury cases (+) and controls (−) from canine whole blood. Columns 2-4 represent replicates of mitochondrial gene copy number, and columns 6-8 represent replicates of B2M copy number. Mitochondrial DNA content (Mt/N ratio) is shown in column 10.

| Brain injury | Mito1 | Mito2 | Mito3 | Average | B2M1 | B2M2 | B2M3 | Average | Mt/N |
| --- | --- | --- | --- | --- | --- | --- | --- | --- | --- |
| - | 348999 | 330703 | 342022 | 340575 | 6759 | 7135 | 6992 | 6962 | 98 |
| - | 1098783 | 1121188 | 1128758 | 1116243 | 8333 | 7630 | 7326 | 7763 | 288 |
| - | 654481 | 624369 | NA | 639425 | 6845 | 7376 | NA | 7110 | 180 |
| - | 733797 | 743738 | NA | 738768 | 7178 | 6845 | NA | 7012 | 211 |
| - | 643914 | 639594 | 648263 | 643924 | 9933 | 10343 | 9474 | 9917 | 130 |
| - | 536913 | 488630 | 505354 | 510299 | 4955 | 1874 | 5337 | 4055 | 252 |
| + | 562815 | 547864 | 544189 | 551623 | 5301 | 4758 | 4447 | 4835 | 228 |
| + | 309177 | 344332 | 319759 | 324423 | 3238 | 3441 | 3992 | 3557 | 182 |
| + | 461253 | 392468 | 440031 | 431251 | 5114 | 5011 | 5778 | 5301 | 163 |
| + | 1106201 | 1300075 | 1447854 | 1284710 | 10566 | 11938 | NA | 11252 | 228 |
| + | 748760 | 795505 | 774379 | 772881 | 7326 | 7476 | NA | 7401 | 209 |
| + | 868225 | 758904 | NA | 813565 | 3618 | 3569 | NA | 3594 | 453 |

Even though the values were higher and had a larger spread, blood mtDNA-CN in the dogs with and without injury did not significantly differ (Figure 4).

**Figure 4.**
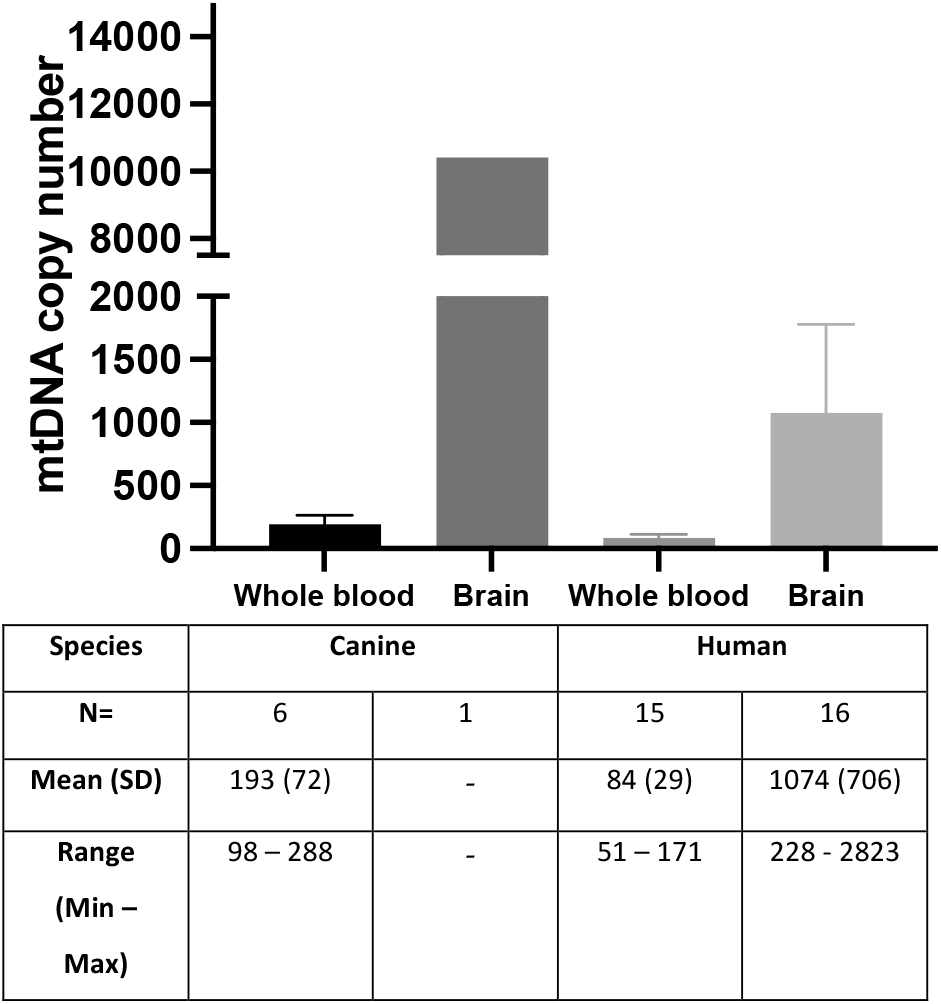
MtDNA-CN in the whole blood and in the brain of dogs and humans. DNA was extracted from whole blood and brain samples collected from dogs and individuals without any brain injury. MtDNA-CN was assessed as a ratio of mitochondrial genome to nuclear genome (Mt/N*2) using real-time qPCR. Data shown as mean ± standard deviation. Human data obtained from research previously published by our group [18].

**Figure 5.**
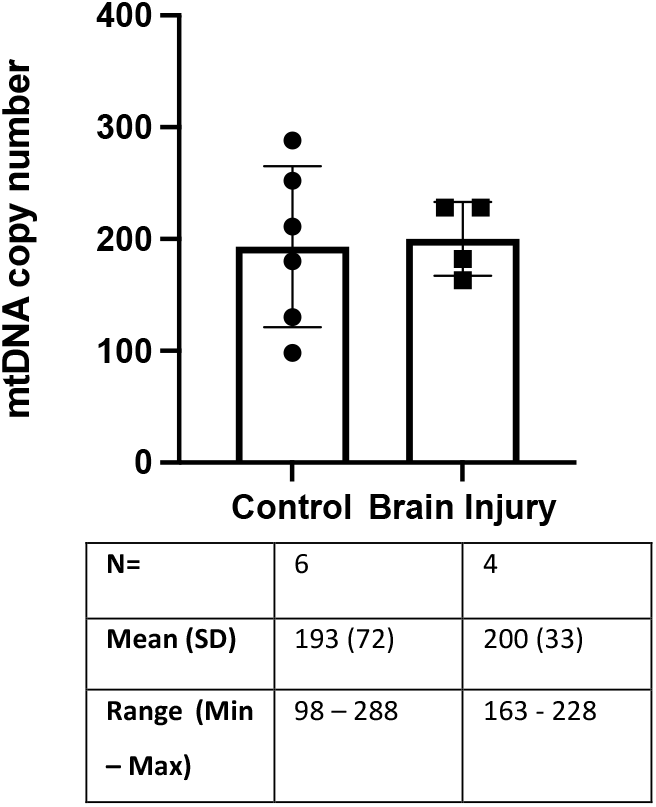
MtDNA-CN in the whole blood of dogs with brain injury and controls. Whole blood samples were collected from canines with brain injury and control. Following DNA extraction and template pre-treatment, MtDNA-CN was assessed as a ratio of mitochondrial genome to nuclear genome (Mt/N*2) using real-time qPCR. Data shown as mean ± standard deviation, n=4-6.

In the cerebral cortex of a single dog with brain injury, mtDNA-CN was 8650 copies/genome, while in the control dog it was 10417 copies/genome (Figure 4).

### 3.4 Time-dependent changes in brain mtDNA-CN following brain injury

We next used the assay to monitor blood mtDNA-CN over 5 days in a single dog that experienced a global hypoxic-ischaemic event caused by cardiopulmonary arrest during general anaesthesia. Three blood samples were collected between days 1 and 5. Twelve hours post-injury, mtDNA-CN was 288 copies/genome, declining to 209 copies/genome on day 3. By day 5, mtDNA-CN increased to 453 copies/genome, which exceeded the range observed in control blood samples (Figure 6).

**Figure 6.**
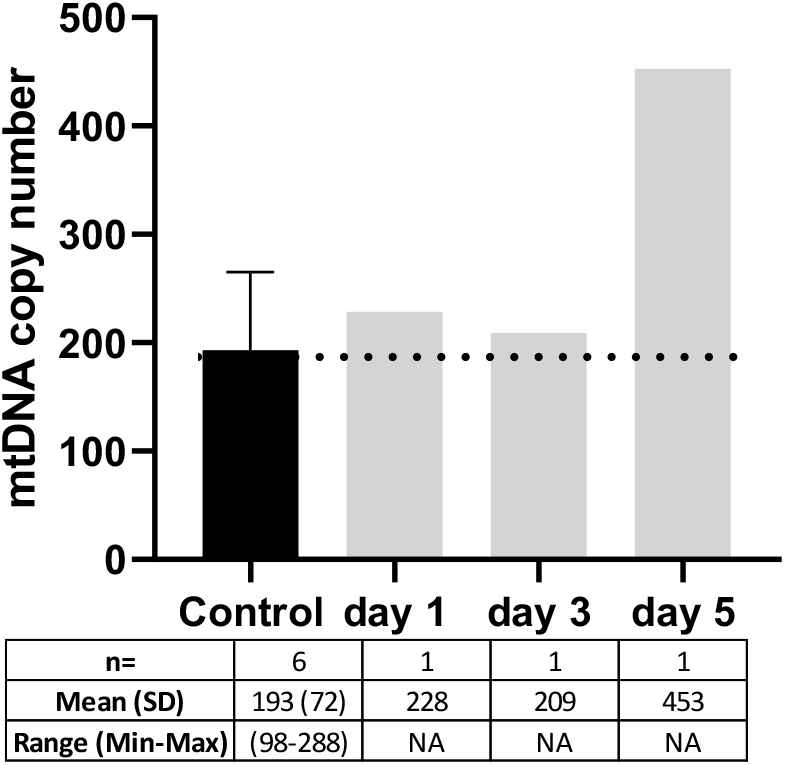
Longitudinal study of blood mtDNA-CN in a 5-year-old female neutered Shih Tzu dog following a global hypoxic-ischemic brain injury. Three blood samples were collected from a single animal following hypoxic-ischaemic brain injury. DNA was extracted using a standard column-based technique and mtDNA-CN was assessed as the ratio of mitochondrial genome to nuclear genome (Mt/N*) using real-time qPCR. Data shown as absolute levels of mtDNA-CN, n=1/day. Control = canine blood samples without any brain injury.

## 4. Discussion

Brain injury in dogs can have severe long-term consequences, but limited tools exist to support diagnosis and prognostication. While advanced imaging is valuable in lesion identification and characterisation, it is costly and requires sedation or general anaesthesia. A serum biomarker to non-invasively and cost effectively support diagnosis and prognostication would be highly valuable in veterinary medicine[29].

MtDNA content has been shown to be altered in tissues and in circulating cells in a wide variety of diseases in human studies,, including diabetes [30], multiple sclerosis [31] and cancer [32] and has been proposed as a biomarker of mitochondrial dysfunction [9]. MtDNA content changes in the peripheral blood have been previously reported in a porcine model, where an increase was detected 6 and 25 hours after traumatic brain injury, correlating with changes in mitochondrial respiration [20]. Following brain injury, brain mitochondria are subjected to increased levels of stressors including excitatory neurotransmitters, Ca^2+^ and reactive oxygen species, leading to mitochondrial dysfunction, mtDNA damage and mutagenesis, eventually resulting in neuronal death [19]. Given the lack of a suitable clinical biomarker and prior evidence of mitochondrial dysfunction in brain injury, we proposed to evaluate the use of circulating mtDNA-CN as a possible biomarker of canine acute brain injury in veterinary medicine. To test this idea, we designed a specific qPCR assay which can accurately measure canine mtDNA-CN. We identified unique regions in the canine mtDNA genome which were not duplicated in the form of NumtS, and these were used to design oligonucleotide primers for the absolute quantification assay using real time qPCR. We determined mtDNA-CN in canine blood and brain samples, and we evaluated the potential of measuring circulating mtDNA-CN as a minimally invasive blood test to detect canine brain injury.

As previously reported for many species including human [12] and mouse [24], we found that sequences virtually identical to sections spanning more than 95% of the mitochondrial genome are present in multiple chromosomal regions in the canine nuclear genome in the form of NumtS. The presence of NumtS should be taken into account when measuring mtDNA-CN to avoid [33]. We detected NumtS in 30 out of 39 chromosomes, varying from 34-3944bp, with the largest being identical to ∼36% of the entire mtDNA sequence. Our observation of multiple NumtS in the canine genome is in line with recent reports by *Edwards et al*., *2021* who identified nearly 300 Numts in the Basenji dog; these NumtS spanned nearly the whole canine mitochondrial genome [34]. It was important in our assay design to ensure that the primers we used do not co-amplify NumtS. To achieve this, we identified a unique region in the mitochondrial genome that matched both the Boxer and Golder Retriever, two dog breeds with dissimilarities between their mitochondrial genomes. This unique region was identified between position 14,231-14,403 and 6,927-7,099 bp respectively and allowed us to design mitochondrial primers without co-amplification of NumtS. We found that canine blood contains from 98 to 288 mtDNA copies/genome with a mean of 193 ±72. These levels are similar to those previously reported in human blood samples [35-37] and in mice, where levels ranging from 80 to 344 copies/genome have been reported [24]. As in humans, variations in the mitochondrial genome have been reported in the domestic dog and specific mitochondrial defects have been reported in some studies reviewed by Tkaczyk-Wlizło *et al*., 1998 [38]. Although mtDNA-CN changes are proposed as a potential mechanism leading to mitochondrial dysfunction, only a few studies have measured mtDNA-CN in dogs, all using relative quantification [26, 27]. The assay described here enables absolute quantification, allowing direct comparisons between tissues and could be applied to studies of mitochondrial disease in dogs, with translational potential to human mitochondrial diseases.

We performed analysis of serial blood samples in a 5-year-old, female neutered Shih Tzu dog. The samples were collected 12 hours, 3 days and 5 days after a cardiopulmonary arrest under general anaesthesia. The dog had pronounced neurological deficits following resuscitation, consistent with a global hypoxic-ischaemic brain injury. Interestingly, mtDNA-CN was most elevated at 5 days, suggesting a potential lag between insult and release of mtDNA from injured brain cells. The mtDNA-CN was also the highest measured across all analysed samples in our study, suggesting that a severe, diffuse injury such as a global hypoxic-ischemic insult might cause widespread neuronal compromise and subsequent mtDNA release. The dog ultimately went on to make a good neurological recovery and future studies are warranted to evaluate if the extent and the duration of the elevation in plasma mtDNA-CN correlate to prognosis.

We analysed mtDNA-CN in cerebral cortex and, as expected, found the levels to be higher compared with those found in blood (∼10 fold higher), though the analysis was limited to a single dog. While we compared a dog with a normal brain and a dog with “brain injury”, the latter had a resolving focal cerebral haemorrhagic event (secondary to a suspected coagulopathy) and hence the small sample of brain we analysed might not be representative of the lesion. Future studies comparing healthy canine cerebral cortex with cerebral cortex from a range of lesions are needed.

Our study has several limitations. Our sample sizes were small and heterogenous in terms of dog age, breed, sex and underlying disorder. The dogs we used as controls, whilst there was no evidence of brain injury, were cases that presented to a referral veterinary hospital and hence had a range of presenting complaints, some of which might have altered the blood mtDNA-CN. Therefore, we cannot be sure that the values obtained from our control animal data is reflective of healthy dogs. Furthermore, for the samples from dogs with brain injury, the underlying diagnoses varied, as did the severity of their presentations. Some dogs had focal brain injury, whilst others had a much more diffuse encephalopathy. Furthermore, the precise timing of injury/disease onset relative to blood sampling is variable and undefined, and if the mtDNA-CN in blood showed an initial acute rise and then a progressive decline, the time of sampling would significantly affect the data produced. This variability could lead to large variations in the data. Overall, our findings provide descriptive data and an initial exploratory comparison between dogs with and without brain injury but further studies are required to fully evaluate its potential clinical efficacy.

## 5. Conclusions

In conclusion, we have shown that it is possible to accurately quantify mtDNA-CN in canine blood samples, and our data suggests that these levels may change in brain injury. A biomarker for canine brain injury could provide a sensitive, cost-effective, and easily accessible tool for both diagnosis and prognostication in veterinary medicine. Therefore, we propose that this assay be utilized to determine its usefulness as a clinical biomarker of brain injury.

## Conflict of interest

Authors do not have competing conflict of interest.

## Notes

### Competing Interest Statement

The authors have declared no competing interest.

